# A draft genome assembly of the eastern banjo frog *Limnodynastes dumerilii dumerilii* (Anura: Limnodynastidae)

**DOI:** 10.1101/2020.03.03.971721

**Authors:** Qiye Li, Qunfei Guo, Yang Zhou, Huishuang Tan, Terry Bertozzi, Yuanzhen Zhu, Ji Li, Stephen Donnellan, Guojie Zhang

## Abstract

Amphibian genomes are usually challenging to assemble due to large genome size and high repeat content. The Limnodynastidae is a family of frogs native to Australia, Tasmania and New Guinea. As an anuran lineage that successfully diversified on the Australian continent, it represents an important lineage in the amphibian tree of life but lacks reference genomes. Here we sequenced and annotated the genome of the eastern banjo frog *Limnodynastes dumerilii dumerilii* to fill this gap. The total length of the genome assembly is 2.38 Gb with a scaffold N50 of 285.9 kb. We identified 1.21 Gb of non-redundant sequences as repetitive elements and annotated 24,548 protein-coding genes in the assembly. BUSCO assessment indicated that more than 94% of the expected vertebrate genes were present in the genome assembly and the gene set. We anticipate that this annotated genome assembly will advance the future study of anuran phylogeny and amphibian genome evolution.

## Introduction

The recent powerful advances in genome sequencing technology have allowed efficient decoding of the genomes of many species [1, 2]. So far, genome sequences are available publicly for more than one thousand species sampled across the animal branch of the tree of life. These genomic resources have provided vastly improved perspectives on our knowledge of the origin and evolutionary history of metazoans [3, 4], facilitated advances in agriculture [5], enhanced approaches for conservation of endangered species [6], and uncovered the genomic changes underlying the evolutionary successes of some clades such as birds [7] and insects [8]. However, amphibian genomes are still challenging to assemble due to their large genome sizes, high repeat content and sometimes high heterozygosity if specimens are collected from wild populations [9]. This also accounts for the scarcity of reference genomes for Anura (frogs and toads) — the most species-rich order of amphibians including many important models for developmental biology and environmental monitoring [10]. Specifically, despite the existence of more than 7,000 living species of Anura [11], only 10 species have their genomes sequenced and annotated to date [12–21], which cover only 8 out of the 54 anuran families. Moreover, genomes of Neobatrachia, which contains more than 95% of the anuran species [11], are particularly under-represented. Only 5 of the 10 publicly available anuran genomes belong to Neobatrachia [22]. This deficiency of neobatrachian genomes would undoubtedly restrict the study of the genetic basis underlying the great diversification of this amphibian lineage, and our understanding of the adaptive genomic changes that facilitate the aquatic to terrestrial transition of vertebrates and the numerous unique reproductive modes found in this clade.

As a candidate species proposed for genomic analysis by the Genome 10K (G10K) initiative [9], we sequenced and annotated the genome of the Australian banjo frog *Limnodynastes dumerilii* (also called the pobblebonk; NCBI:txid104065) to serve as a representative species of the neobatrachian family Limnodynastidae. This burrowing frog is endemic to Australia and named after its distinctive “bonk” call, which is likened to a banjo string being plucked. It mainly occurs along the southeast coast of Australia, from the coast of New South Wales, throughout Victoria and into the southwest corner of South Australia and Tasmania [23]. Five subspecies of *L. dumerilii* are recognized, including *Limnodynastes dumerilii dumerilii, L. dumerilii grayi, L. dumerilii fryi*, *L. dumerilii insularis* and *L. dumerilii variegata* [24]. The subspecies chosen for sequencing is the eastern banjo frog *L. dumerilii dumerilii* (NCBI:txid104066), as it is the most widespread among the five subspecies and forms hybrid zones with a number of the other subspecies [23]. We believe that the release of genomic resources from this neobatrachian frog will benefit the future studies of phylogenomics and comparative genomics of anurans, and also facilitate other research related to the evolutionary biology of *Limnodynastes*.

## Methods

### Sample collection, library construction and sequencing

Genomic DNA was extracted from the liver of an adult female *Limnodynastes dumerilii dumerilii* (Fig. 1) using the Gentra Puregene Tissue Kit (QIAGEN, Hilden, Germany) according to manufacturer’s instructions with the following exceptions: following the DNA precipitation step, DNA was spooled onto a glass rod, washed twice in 70% ethanol and dried before dissolving in 100 ul of the recommended elution buffer [25]. The specimen was originally caught in River Torrens, Adelaide, South Australia, Australia, and is archived in the South Australian Museum (registration number: SAMAR66870).

**Figure 1.**
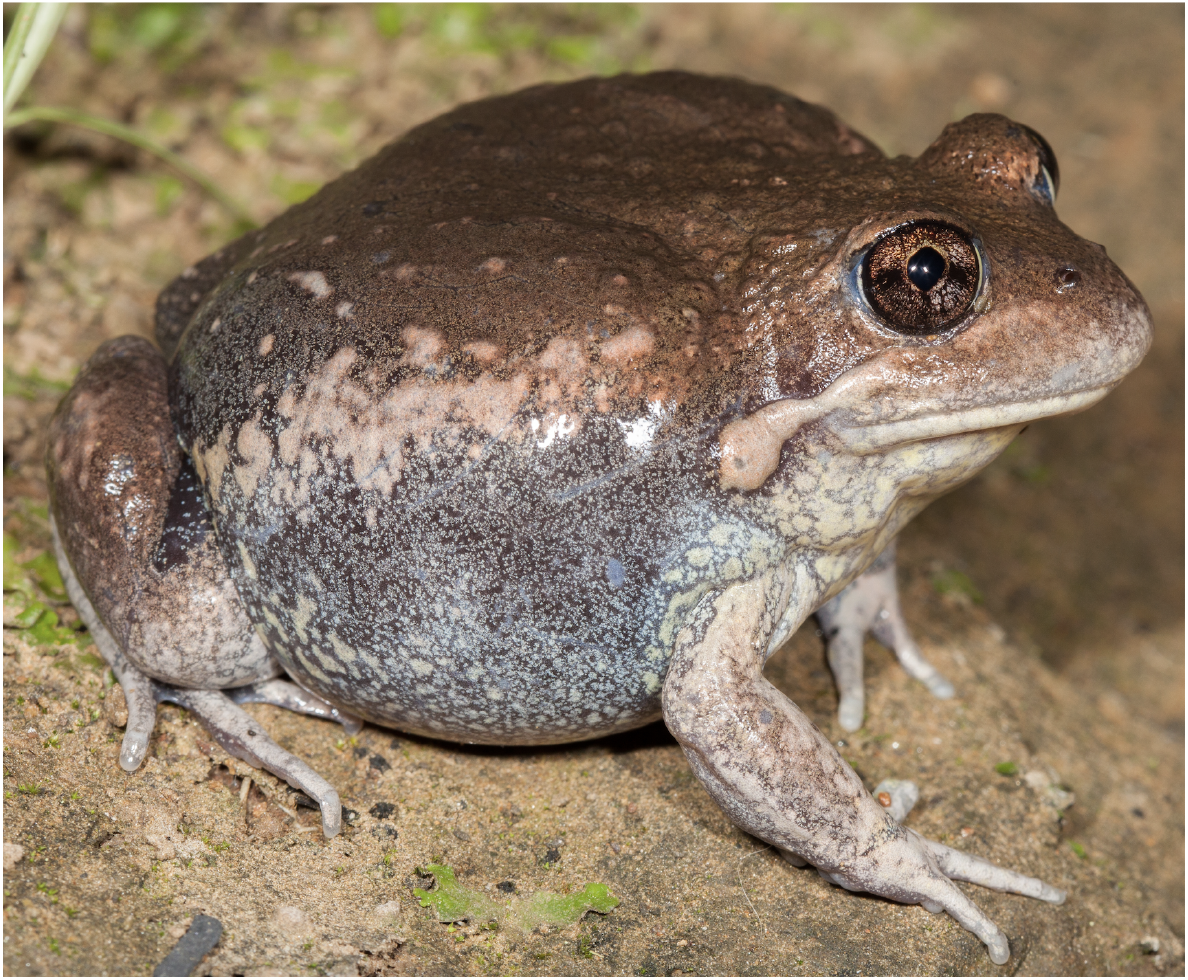
Photograph of an adult *Limnodynastes dumerilii dumerilii* from the Adelaide region (image from Stephen Mahony).

A total of 211 Gb of sequences were generated from four short-insert libraries (170 bp × 1, 250 bp × 1, 500 bp × 1, and 800 bp × 1), and 185 Gb of sequences from ten mate-paired libraries (2 kb × 3, 5 kb × 3, 10 kb × 2, and 20 kb × 2). All the 14 libraries were subjected to paired-end sequencing on the HiSeq 2000 platform following the manufacturer’s instructions (Illumina, San Diego, CA, USA), using PE100 or PE150 chemistry for the short-insert libraries and PE49 for the mate-paired libraries [26] (Table 1).

**Table 1.** Statistics of DNA reads produced for the *L. d. dumerilii* genome.

| NCBI accession | CNSA accession | Library insert size (bp) | Read length (bp) | Raw data |  |  | Clean data |  |  |
| --- | --- | --- | --- | --- | --- | --- | --- | --- | --- |
|  |  |  |  | Total bases (Gb) | Sequence depth (X) | Physical depth (X) | Total bases (Gb) | Sequence depth (X) | Physical depth (X) |
| SRR10802019 | CNR0165422 | 170 | 100 | 43.45 | 17.11 | 14.54 | 36.52 | 14.38 | 12.11 |
| SRR10802018 | CNR0165423 | 250 | 150 | 67.56 | 26.60 | 22.17 | 45.71 | 18.00 | 16.00 |
| SRR10802013 | CNR0165424 | 500 | 150 | 61.47 | 24.20 | 40.33 | 29.79 | 11.73 | 26.12 |
| SRR10802012 | CNR0165425 | 800 | 150 | 38.34 | 15.10 | 40.26 | 18.56 | 7.31 | 21.38 |
| SRR10802011 | CNR0165426 | 2,000 | 49 | 18.79 | 7.40 | 151.00 | 9.84 | 3.87 | 99.33 |
| SRR10802009 | CNR0165427 | 2,000 | 49 | 19.86 | 7.82 | 159.53 | 8.70 | 3.43 | 87.84 |
| SRR10802008 | CNR0165428 | 2,000 | 49 | 21.25 | 8.36 | 170.71 | 10.38 | 4.09 | 104.75 |
| SRR10802007 | CNR0165429 | 5,000 | 49 | 18.60 | 7.32 | 373.70 | 3.92 | 1.54 | 98.94 |
| SRR10802010 | CNR0165430 | 5,000 | 49 | 18.03 | 7.10 | 362.19 | 3.46 | 1.36 | 87.39 |
| SRR10802006 | CNR0165431 | 5,000 | 49 | 15.47 | 6.09 | 310.78 | 1.87 | 0.74 | 47.25 |
| SRR10802017 | CNR0165432 | 10,000 | 49 | 16.07 | 6.33 | 645.68 | 1.45 | 0.57 | 73.13 |
| SRR10802016 | CNR0165433 | 10,000 | 49 | 20.74 | 8.17 | 833.24 | 3.45 | 1.36 | 174.07 |
| SRR10802015 | CNR0165434 | 20,000 | 49 | 16.93 | 6.66 | 1360.12 | 0.98 | 0.38 | 98.44 |
| SRR10802014 | CNR0165435 | 20,000 | 49 | 19.09 | 7.52 | 1533.74 | 1.44 | 0.57 | 145.78 |
| Total |  |  |  | 395.66 | 155.77 | 6018.00 | 176.07 | 69.32 | 1092.54 |

Note: Depth calculation was based on the estimated haploid genome size of 2.54 Gb according to *k*-mer analysis. Sequence depth is the average number of times a base is read, while physical depth is the average number of times a base is spanned by sequenced DNA fragments.

The raw sequencing data from each library were subjected to strict quality control by SOAPnuke (v1.5.3, RRID:SCR_015025) [27] prior to downstream analyses (see [28] for detailed parameters for each library). Briefly, for the raw reads from each library, we trimmed the unreliable bases at the head and tail of each read where the per-position GC content was unbalanced or the per-position base quality was low across all reads; we removed the read pairs with adapter contamination, with high proportion of low-quality or unknown (N) bases; we removed duplicate read pairs potentially resulted from polymerase chain reaction (PCR) amplification (i.e. PCR duplicates); and we also removed the overlapping read pairs in all but the 170 bp and 250 bp libraries where the paired reads were expected to be overlapping. As shown in Table 2, data reduction in the short-insert libraries were mainly caused by the truncation of the head and tail of each read and the discard of read pairs with too many low-quality bases. But it is noteworthy that PCR duplication rates for all the short-insert libraries are extremely low (0.2% – 2.6%), indicating that sequences from these libraries are diverse. In contrast, data reduction in the mate-paired libraries were mainly due to the discard of PCR duplicates, which made up 22.6% – 83.0% of the raw data (Table 2). A total of 176 Gb of clean sequences were retained for genome assembly after these strict quality controls, representing 69 times coverage of the estimated haploid genome size of *L. d. dumerilii* in terms of sequence depth, and 1,093 times in terms of physical depth (Table 1).

**Table 2.** The summary of data filtering for each library.

| NCBI accession | CNSA accession | Library insert size (bp) | % Discarded bases | % of bases discarded due to different factors |  |  |  |  |  |
| --- | --- | --- | --- | --- | --- | --- | --- | --- | --- |
|  |  |  |  | Adapter contamination (-f & -r) | Low quality bases (-l & -q) | N bases (-n) | Small insert size (-s) | PCR duplicates (-d) | Trimming (-t) |
| SRR10802019 | CNR0165422 | 170 | 15.95 | 0.18 | 8.36 | 0.38 | 0.00 | 2.62 | 4.42 |
| SRR10802018 | CNR0165423 | 250 | 32.34 | 0.22 | 23.66 | 0.13 | 0.00 | 0.81 | 7.52 |
| SRR10802013 | CNR0165424 | 500 | 51.54 | 0.18 | 26.42 | 0.14 | 6.65 | 0.52 | 17.62 |
| SRR10802012 | CNR0165425 | 800 | 51.59 | 0.05 | 39.25 | 0.62 | 6.15 | 0.15 | 5.38 |
| SRR10802011 | CNR0165426 | 2,000 | 47.64 | 0.28 | 4.51 | 0.32 | 6.48 | 22.63 | 13.43 |
| SRR10802009 | CNR0165427 | 2,000 | 56.18 | 0.16 | 4.58 | 0.18 | 5.75 | 34.27 | 11.24 |
| SRR10802008 | CNR0165428 | 2,000 | 51.16 | 0.13 | 5.36 | 0.20 | 5.59 | 27.36 | 12.52 |
| SRR10802007 | CNR0165429 | 5,000 | 78.93 | 0.08 | 4.47 | 0.17 | 3.11 | 65.69 | 5.40 |
| SRR10802010 | CNR0165430 | 5,000 | 80.80 | 0.78 | 2.84 | 0.83 | 3.03 | 68.38 | 4.92 |
| SRR10802006 | CNR0165431 | 5,000 | 87.90 | 8.45 | 2.44 | 0.73 | 2.27 | 70.89 | 3.10 |
| SRR10802017 | CNR0165432 | 10,000 | 90.99 | 0.23 | 4.23 | 0.12 | 2.89 | 81.20 | 2.31 |
| SRR10802016 | CNR0165433 | 10,000 | 83.37 | 3.95 | 6.35 | 0.18 | 2.29 | 66.35 | 4.26 |
| SRR10802015 | CNR0165434 | 20,000 | 94.24 | 0.62 | 3.71 | 0.10 | 5.29 | 83.04 | 1.48 |
| SRR10802014 | CNR0165435 | 20,000 | 92.44 | 1.11 | 5.44 | 0.68 | 3.90 | 79.37 | 1.94 |

Note: The options of SOAPnuke (v1.5.3) that control the corresponding factors are indicated in parentheses. The detailed settings of these options for each library are deposited at protocols.io [28].

### Genome size estimation and genome assembly

To obtain a robust estimation of the genome size of *L. d. dumerilii*, we conducted *k*-mer analysis with all of the clean sequences (131 Gb) from the four short-insert libraries using a range of *k* values (17, 19, 21, 23, 25, 27, 29 and 31). The *k*-mer frequencies were counted by Jellyfish (v2.2.6) [29] with the *-C* setting. The genome size of *L. d. dumerilii* was estimated to be around 2.54 Gb (Table 3), which was calculated as the number of effective *k*-mers (i.e. total *k*-mers – erroneous *k*-mers) divided by the homozygous peak depth following Cai *et al* [30]. It is noteworthy that, the presence of a distinct heterozygous peak, which displayed half of the depth of the homozygous peak in the *k*-mer frequency distribution, suggests that the diploid genome of this wild-caught individual has a high level of heterozygosity (Fig. 2). The rate of heterozygosity was estimated to be around 1.17% by GenomeScope (v1.0.0, RRID:SCR_017014) [31] (Table 3).

**Table 3.**
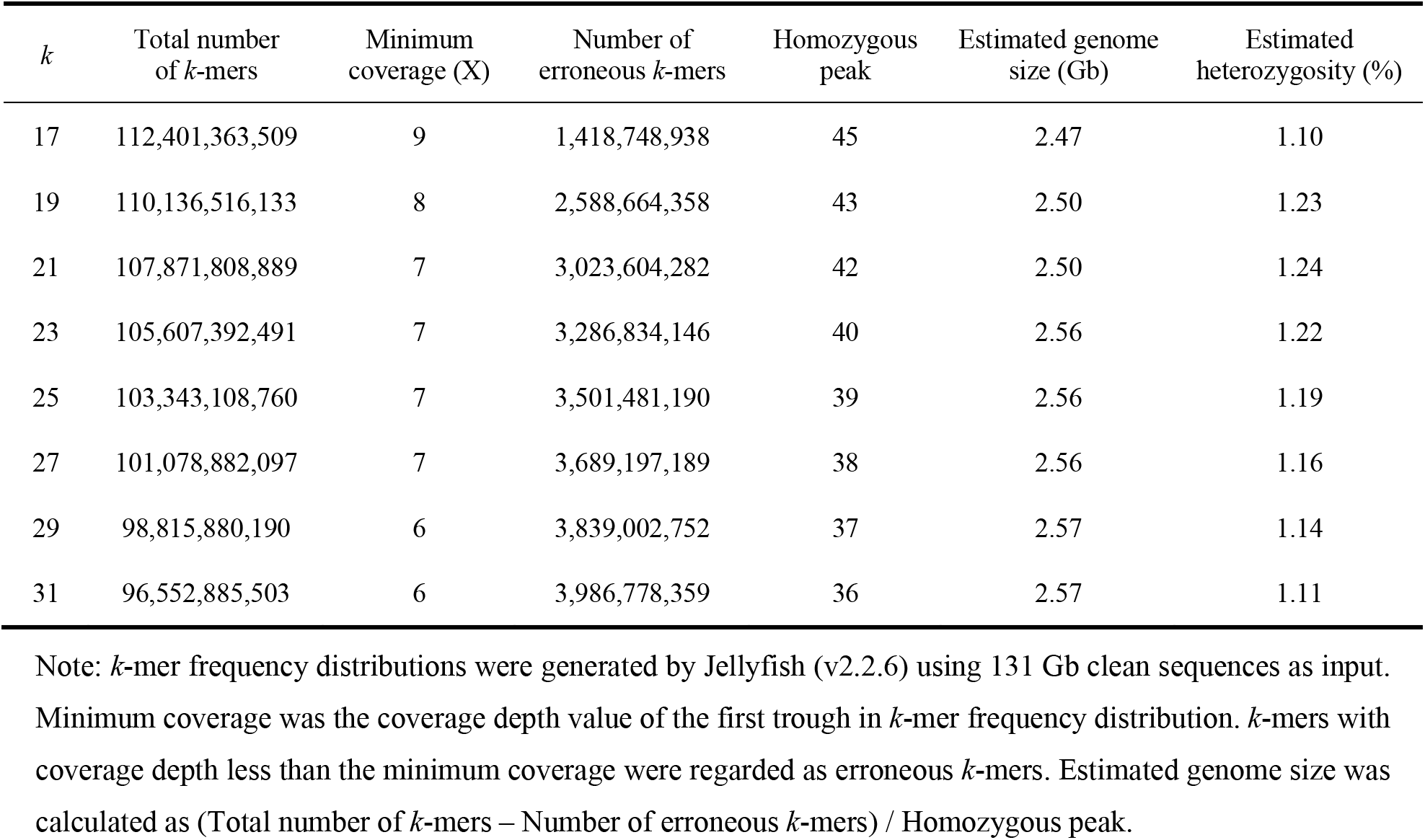
Estimation of genome size and heterozygosity of *L. d. dumerilii* by *k*-mer analysis.

**Figure 2.**
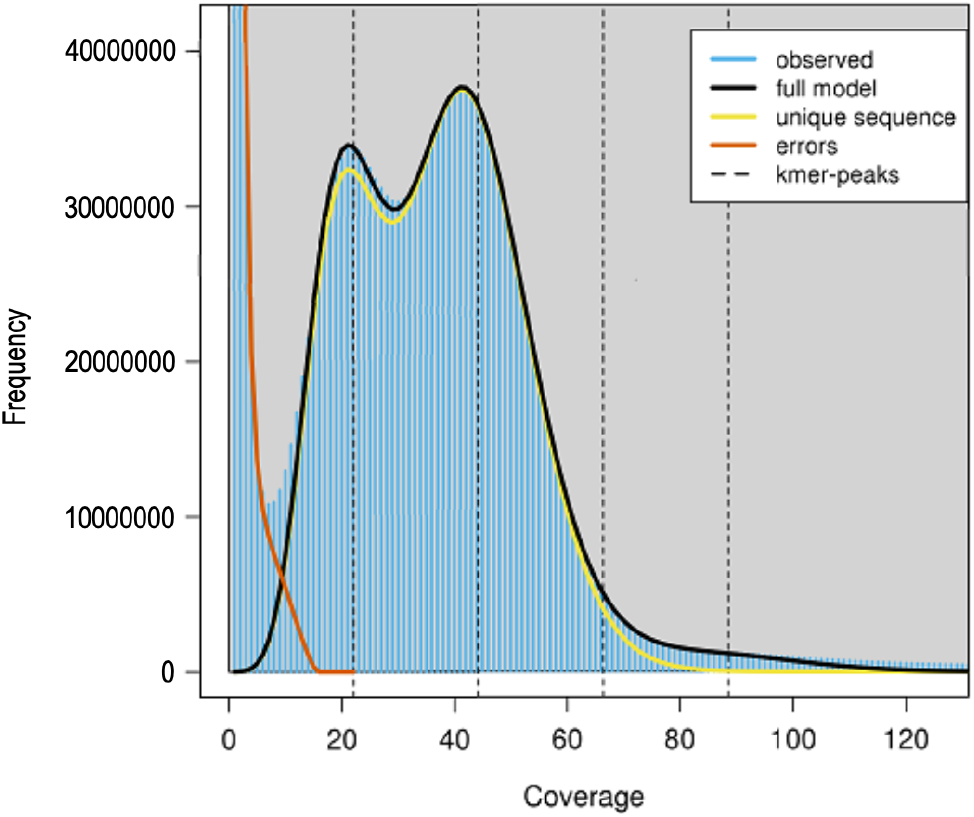
A 21-mer frequency distribution of the *L. d. dumerilii* genome data. The first peak at coverage 21X corresponds to the heterozygous peak. The second peak at coverage 42X corresponds to the homozygous peak.

We then employed Platanus (v1.2.1, RRID:SCR_015531) [32] to assemble the genome of *L. d. dumerilii*. Briefly, all the clean sequences from the four short-insert libraries were first assembled into contigs using *platanus assemble* with parameters *-t 20 -k 29 -u 0.2 -d 0.6 -m 150*. Then paired-end reads from the four short-insert and ten mate-paired libraries were used to connect contigs into scaffolds by *platanus scaffold* with parameters *-t 20 -u 0.2 -l 3* and the insert size information of each library. Finally, *platanus gap_close* was employed to close intra-scaffold gaps using the paired-end reads from the four short-insert libraries with default settings. This Platanus assembly was further improved by Kgf (version 1.16) [9] followed by GapCloser (v1.10.1, RRID:SCR_015026) [9] for gap filling with the clean reads from the four short-insert libraries.

### Repetitive element annotation

Both homology-based and *de novo* predictions were employed to identify repetitive elements in the *L. d. dumerilii* genome assembly [33]. For homology-based prediction, known repetitive elements were identified by aligning the *L. d. dumerilii* genome sequences against the Repbase-derived RepeatMasker libraries using RepeatMasker (v4.1.0, RRID:SCR_012954; setting - *nolow -norna -no_is*) [34], and against the transposable element protein database using RepeatProteinMask (an application within the RepeatMasker package; setting *-noLowSimple - pvalue 0.0001 -engine ncbi*). For *de novo* prediction, RepeatModeler (v2.0, RRID:SCR_015027) [35] was first executed on the *L. d. dumerilii* assembly to build a *de novo* repeat library for this species. Then RepeatMasker was employed to align the *L. d. dumerilii* genome sequences against the *de novo* library for repetitive element identification. Tandem repeats in the *L. d. dumerilii* genome assembly were identified by Tandem Repeats Finder (v4.09) [36] with parameters *Match=2 Mismatch=7 Delta=7 PM=80 PI=10 Minscore=50 MaxPeriod=2000*.

### Protein-coding gene annotation

Similar to repetitive element annotation, both homology-based and *de novo* predictions were employed to build gene models for the *L. d. dumerilii* genome assembly [37]. For homology-based prediction, protein sequences from diverse vertebrate species (see [37] for the sources), including *Danio rerio, Xenopus tropicalis, Xenopus laevis, Nanorana parkeri, Microcaecilia unicolor, Rhinatrema bivittatum, Anolis carolinensis, Gallus gallus* and *Homo sapiens*, were first aligned to the *L. d. dumerilii* genome assembly using TBLASTN (blast-2.2.26, RRID:SCR_011822) [38] with parameters *-F F -e 1e-5*. Then the genomic sequences of the candidate loci together with 5 kb flanking sequences were extracted for exon-intron structure determination, by aligning the homologous proteins to these extracted genomic sequences using GeneWise (wise-2.2.0, RRID:SCR_015054) [39]. For *de novo* prediction, we randomly picked 1,000 homology-derived gene models of *L. d. dumerilii* with complete open reading frames (ORFs) and reciprocal aligning rates exceeding 90% against the *X. tropicalis* proteins to train AUGUSTUS (v3.3.1, RRID:SCR_008417) [40]. The obtained gene parameters were then used by AUGUSTUS to predict protein-coding genes on the repeat-masked *L. d. dumerilii* genome assembly. Finally, gene models derived from the above two methods were combined into a non-redundant gene set using a similar strategy to Xiong *et al*. (2016) [41]. Genes showing BLASTP (blast-2.2.26, RRID:SCR_001010; parameters *-F F -e 1e-5*) hits to transposon proteins in the UniProtKB/Swiss-Prot database (v2019_11), or with more than 70% of their coding regions overlapping repetitive sequences, were removed from the combined gene set.

## Results and Discussion

### Assembly and annotation of the *L. d. dumerilii* genome

We assembled the nuclear genome of a female eastern banjo frog *L. d. dumerilii* (Fig. 1) with ~176 Gb (69X) clean Hiseq data from four short-insert libraries (170 bp × 1, 250 bp × 1, 500 bp × 1, and 800 bp × 1) and ten mate-paired libraries (2 kb × 3, 5 kb × 3, 10 kb × 2, and 20 kb × 2) (Table 1–2). The final genome assembly comprised 520,896 sequences with contig and scaffold N50s of 10.2 kb and 286.0 kb, respectively, and a total length of 2.38 Gb, which is close to the estimated genome size of 2.54 Gb by *k*-mer analysis (Table 3–4 and Fig. 2). There are 242 Mb of regions present as unclosed gaps (Ns), accounting for 10.2% of the assembly. The GC content of the *L. d. dumerilii* assembly excluding gaps was estimated to be 41.0% (Table 4). The combination of homology-based and *de novo* prediction methods masked 1.21 Gb of non-redundant sequences as repetitive elements, accounting for 56.4 % of the *L. d. dumerilii* genome assembly excluding gaps (Table 5). We also obtained 24,548 protein-coding genes in the genome assembly, of which 67% had complete ORF. Functional annotation by searching the *L. d. dumerilii* proteins against public databases of UniProtKB/Swiss-Prot (v2019_11, RRID:SCR_004426) [42], NCBI nr (v20191030), and KEGG (v93.0, RRID:SCR_012773) [43] with BLASTP (blast-2.2.26; parameters *-F F -e 1e-5*) successfully annotated almost all of the *L. d. dumerilii* gene loci (Table 6).

**Table 4.**
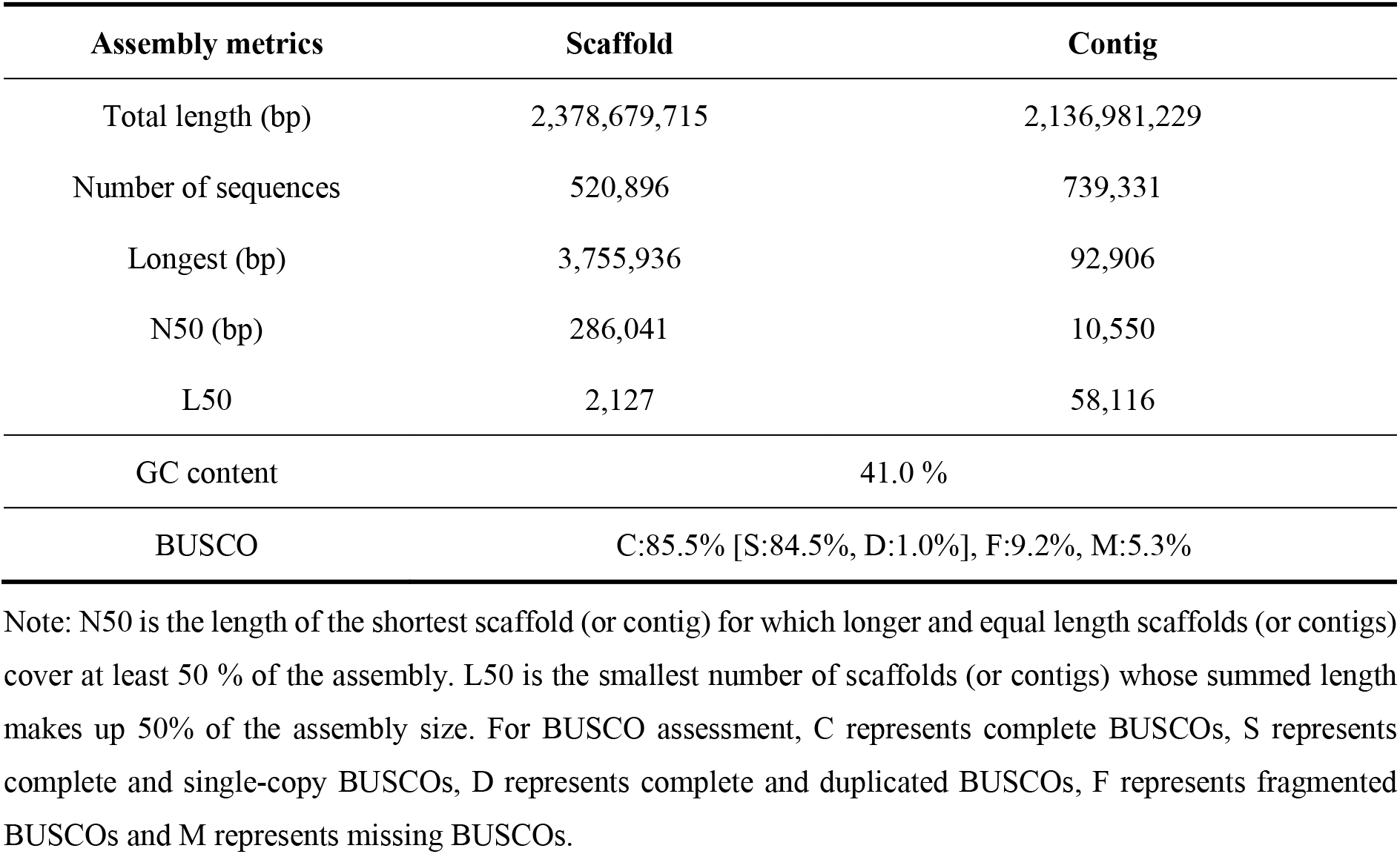
Metrics for the *L. d. dumerilii* genome assembly.

| Assembly metrics | Scaffold | Contig |
| --- | --- | --- |
| Total length (bp) | 2,378,679,715 | 2,136,981,229 |
| Number of sequences | 520,896 | 739,331 |
| Longest (bp) | 3,755,936 | 92,906 |
| N50 (bp) | 286,041 | 10,550 |
| L50 | 2,127 | 58,116 |
| GC content | 41.0 % |  |
| BUSCO | C:85.5% [S:84.5%, D:1.0%], F:9.2%, M:5.3% |  |

Note: N50 is the length of the shortest scaffold (or contig) for which longer and equal length scaffolds (or contigs) cover at least 50 % of the assembly. L50 is the smallest number of scaffolds (or contigs) whose summed length makes up 50% of the assembly size. For BUSCO assessment, C represents complete BUSCOs, S represents complete and single-copy BUSCOs, D represents complete and duplicated BUSCOs, F represents fragmented BUSCOs and M represents missing BUSCOs.

**Table 5.** Statistics of repetitive sequences identified in the *L. d. dumerilii* genome.

| Category | Total repeat length (bp) | % of assembly |
| --- | --- | --- |
| DNA | 155,988,597 | 7.30% |
| LINE | 242,754,702 | 11.36% |
| SINE | 11,761,904 | 0.55% |
| LTR | 97,615,246 | 4.57% |
| Tandem repeats | 178,355,571 | 8.35% |
| Unknown | 704,263,255 | 32.96% |
| Combined | 1,205,873,056 | 56.43% |

Note: DNA: DNA transposon; LINE: long interspersed nuclear element; SINE: short interspersed nuclear elements; LTR: long terminal repeat.

**Table 6.** Summary of protein-coding genes annotated in the *L. d. dumerilii* genome.

| Characteristics of protein-coding genes |  |
| --- | --- |
| Total number of protein-coding genes | 24,548 |
| Gene space (exon + intron; Mb) | 634.6 (26.7 % of assembly) |
| Mean gene size (bp) | 25,851 |
| Mean CDS length (bp) | 1,552 |
| Exon space (Mb) | 38.1 (1.6 % of assembly) |
| Mean exon number per gene | 8.6 |
| Mean exon length (bp) | 181 |
| Mean intron length (bp) | 3,217 |
| Functional annotation by searching public databases |  |
| % of proteins with hits in UniProtKB/Swiss-Prot | 95.8 |
| % of proteins with hits in NCBI nr database | 99.6 |
| % of proteins with KO assigned by KEGG | 71.3 |
| % of proteins with functional annotation (combined) | 99.9 |

### Data validation and quality control

Two strategies were employed to estimate the completeness of the *L. d. dumerilii* genome assembly. First, all the clean reads from the short-insert libraries were aligned to the genome assembly using BWA-MEM (BWA, version 0.7.16, RRID:SCR_010910) with default parameters [44]. We observed that 99.6 % of reads could be mapped back to the assembled genome and 85.6 % of the inputted reads were mapped in proper pairs as accessed by samtools flagstat (SAMtools v1.7, RRID:SCR_002105), suggesting that most sequences of the *L. d. dumerilii* genome were present in the current assembly. Of note, by comparing the genomic distributions of the properly paired reads and the remaining mapped reads in the final assembly, we observed that the reads which could not be mapped in proper pairs tended to locate on the ends of scaffolds, the flanking regions of assembly gaps and the genomic regions annotated as tandem repeats (Table 7), indicating that these regions likely have lower assembly accuracy than other genomic regions. Secondly, we assessed the *L. d. dumerilii* assembly with Benchmarking Universal Single-Copy Orthologs (BUSCO; v3.0.2, RRID:SCR_015008), a software package that can quantitatively measure genome assembly completeness based on evolutionarily informed expectations of gene content [45], and found that up to 94.7 % of the 2,586 expected vertebrate genes were present in the *L. d. dumerilii* assembly. Furthermore, 85.5% and 84.5 % of the expected genes were identified as complete and single-copy genes, respectively (Table 4). This BUSCO assessment further highlighted the comprehensiveness of the current *L. d. dumerilii* genome assembly in terms of gene space.

**Table 7.** The percentages of properly paired reads and other mapped reads locating on different genomic regions.

| Genomic regions | Properly paired reads | % Properly paired reads | Other mapped reads | % Other mapped reads |
| --- | --- | --- | --- | --- |
| Scaffold ends | 256,786 | 6.42% | 653,026 | 16.33% |
| Near assembly gaps | 450,707 | 11.27% | 1,619,089 | 40.48% |
| Exon | 112,389 | 2.81% | 43,808 | 1.10% |
| Intron | 1,011,320 | 25.28% | 570,089 | 14.25% |
| Tandem repeats | 436,761 | 10.92% | 954,934 | 23.87% |
| Other repeats | 2,565,614 | 64.14% | 2,955,171 | 73.88% |

Note: The percentages were estimated based on 4 million reads randomly selected from each of the two read groups. Scaffold ends: 500 bp regions next to the head or tail of each scaffold; Near assembly gaps: 500 bp flanking region of an assembly gap which contains no less than 50 Ns; Tandem repeats: repeats derived from Tandem Repeats Finder; Other repeats: repeats derived from RepeatMasker, RepeatProteinMask and RepeatModeler.

We then evaluated the completeness of the *L. d. dumerilii* protein-coding gene set with BUSCO (v3.0.2) and DOGMA (v3.0, RRID:SCR_015060) [46], a program that measures the completeness of a given transcriptome or proteome based on a core set of conserved domain arrangements (CDAs). BUSCO analysis showed that 97.1 % of the expected vertebrate genes were present in the *L. d. dumerilii* protein-coding gene set with 88.5 % and 84.5% identified as complete and single-copy genes, respectively, close to that estimated for the genome assembly. Meanwhile, DOGMA analysis based on PfamScan Annotations (PfamScan v1.5; Pfam v32.0, RRID:SCR_015060) [47] and the eukaryotic core set identified 95.4 % of the expected CDAs in the annotated gene set. These results demonstrated the high completeness of the *L. d. dumerilii* protein-coding gene set.

### Re-use potential

Here, we report a draft genome assembly of the eastern banjo frog *L. d. dumerilii*. It represents the first genome assembly from the family Limnodynastidae (Anura: Neobatrachia). Although the continuity of the assembly in terms of contig and scaffold N50s is modest, probably due to the high repeat content (56%) and heterozygosity (1.17%), the completeness of this draft assembly is demonstrated to be high according to read mapping and BUSCO assessment. Thus, it is suitable for phylogenomics and comparative genomics analyses with other available anuran genomes or phylogenomic datasets. In particular, the high-quality protein-coding gene set derived from the genome assembly will be useful for deducing orthologous relationships across anuran species or reconstructing the ancestral gene content of anurans. Due to evolutionary importance of *Limnodynastes* frogs in Australia, the genomic resources released in this study will also support further research on the biogeography of speciation, evolution of male advertisement calls, hybrid zone dynamics, and conservation of *Limnodynastes* frogs.

## Availability of supporting data

The raw sequencing reads are deposited in NCBI under the BioProject accession PRJNA597531 and are also deposited in the CNGB Nucleotide Sequence Archive (CNSA) with accession number CNP0000818. The clean reads that passed quality control, the genome assembly, and the protein-coding gene and repeat annotations are deposited in the GigaScience repository (GigaDB) [48]. The genome assembly is also deposited in NCBI under accession number GCA_011038615.1.

## List of abbreviations

BUSCO: Benchmarking Universal Single-Copy Orthologs
G10K: Genome 10
NCBI: National Center for Biotechnology Information
PCR: Polymerase Chain Reaction
ORF: Open Reading Frame
KEGG: Kyoto Encyclopedia of Genes and Genomes
DOGMA: DOmain-based General Measure for transcriptome and proteome quality Assessment
CDA: Conserved Domain Arrangement
CNGB: China National GeneBank
CNSA: CNGB Sequence Archive.

## Author contributions

G.Z. and Q.L. conceived and supervised the study; T.B. and S.D. prepared the DNA samples; Y.Z. and Q.G. performed *k*-mer analysis and genome assembly; Q.G. and J.L. conducted assessment of assembly quality; H.T. performed protein-coding gene annotation; Y.Z. performed repeat annotation; G.Z. and S.D. contributed reagents/materials/analysis tools; Q.L. wrote the manuscript with the inputs from all authors. All authors read and approved the final manuscript.

